# Deviance detection in subthalamic neural population responses to natural stimuli in bats

**DOI:** 10.1101/2023.07.06.547961

**Authors:** Johannes Wetekam, Julio Hechavarría, Luciana López-Jury, Eugenia González-Palomares, Manfred Kössl

**Author notes:** Corresponding authors: Johannes Wetekam, Manfred Kössl.

## Abstract

Deviance detection describes an increase of neural response strength caused by a stimulus with a low probability of occurrence. This ubiquitous phenomenon has been reported for multiple species, from subthalamic areas to auditory cortex. While cortical deviance detection has been well characterised by a range of studies covering neural activity at population level (mismatch negativity, MMN) as well as at cellular level (stimulus-specific adaptation, SSA), subcortical deviance detection has been studied mainly on cellular level in the form of SSA. Here, we aim to bridge this gap by using noninvasively recorded auditory brainstem responses (ABRs) to investigate deviance detection at population level in the lower stations of the auditory system of a hearing specialist: the bat *Carollia perspicillata*. Our present approach uses behaviourally relevant vocalisation stimuli that are closer to the animals’ natural soundscape than artificial stimuli used in previous studies that focussed on subcortical areas. We show that deviance detection in ABRs is significantly stronger for echolocation pulses than for social communication calls or artificial sounds, indicating that subthalamic deviance detection depends on the behavioural meaning of a stimulus. Additionally, complex physical sound features like frequency- and amplitude-modulation affected the strength of deviance detection in the ABR. In summary, our results suggest that at population level, the bat brain can detect different types of deviants already in the brainstem. This shows that subthalamic brain structures exhibit more advanced forms of deviance detection than previously known.

## 1. Introduction

Like all echolocating bats, *Carollia perspicillata* navigates in the dark by emitting stereotypical acoustic pulses and listening to the echoes reflected off objects in its environment. In addition, this bat species has a large variety of social communication calls[1,2], which is a consequence of its social lifestyle, with groups of more than 100 individuals sharing the same roost[3]. This has led to the development of a broad variety of social communication calls. Echolocation pulses and social communication calls differ from each other in their carrier frequencies and durations, with echolocation pulses being higher in frequency and shorter in time (see Fig. 1a for an example echolocation pulse and social communication call). Those two vocalisation types represent fundamentally different behaviours (navigation and social communication) and can alternate in rapid succession for freely behaving bats. This raises a question that has puzzled neuroethologists for years: How does the bat brain process echolocation and social sounds in a fast and energy-efficient way, when they occur in the same acoustic stream? A theoretical model that explains how the brain efficiently deals with the tremendous amount of input it receives is the predictive coding framework and, in relation to this, the ability of deviance detection[4,5]. According to the predictive coding theory, the brain is constantly creating predictions about the incoming stimuli[6]. When the system encounters an unexpected signal, expectations are updated which is represented by a prediction error component in the electrophysiological response. This makes the identification of regularities and deviants in the incoming stream of signals (i.e., deviance detection) crucial for the predictive coding framework. The present study investigates deviance detection to naturally occurring sounds – echolocation pulses and social communication calls – in the bat species *C. perspicillata*. We focussed on studying deviance detection in subthalamic neural populations of the auditory pathway by combining a naturalistic oddball stimulation paradigm (Fig. 1b) with noninvasively recorded auditory brainstem potentials (ABRs). Two experiments were performed: In experiment 1, an echolocation pulse and a social communication call (Fig. 1a) were presented in an oddball paradigm (Fig. 1b). Additionally, by using two control paradigms (the “Many-Standards” (MS) and the 50 % control, Fig. 1c), we aimed to shed light on the possible underlying neural mechanisms responsible for deviance detection, namely deviant enhancement and repetition suppression of the standard response. These neural mechanisms are affected by both controls in different ways, allowing a more detailed characterisation of the effects than by only using an oddball paradigm. In experiment 2, the effect of different acoustic parameters (e.g., carrier frequency and temporal structure) and the behavioural meaning of the auditory input on subthalamic deviance detection was evaluated by performing a cross-comparison of the responses to different stimuli. The stimuli considered ranged from natural vocalisations on one end, to artificially generated vocalisation-mimics, noise bursts that resemble the vocalisations in their frequency range and duration but not in their temporal structure, on the other end (Fig. 1d).

**Figure 1:**
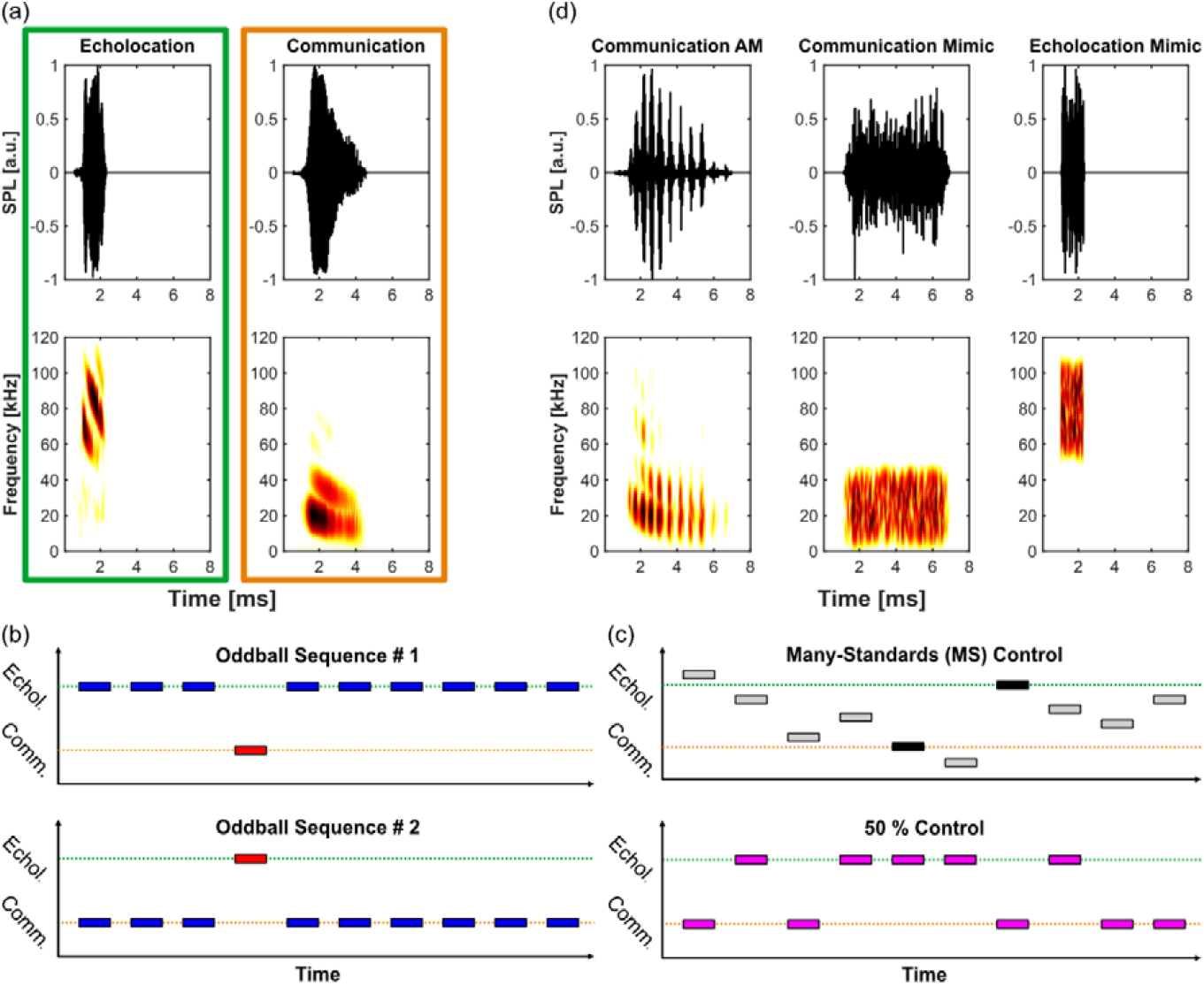
Used stimuli and stimulation protocols. (a) Oscillograms (top) and spectrograms (bottom) of an echolocation pulse (green frame) and a social communication call (orange frame) of C. perspicillata that were used as stimuli in this study. The communication signal is a syllable of a so-called distress call. (b) Schematic representation of the oddball paradigm; blue: standard, red: deviant. (c) Schematic representation of the two control sequences used; Many-Standards control (top) and 50 % control (bottom). (d) Additional stimuli used for a cross-comparison with the vocalisations in (a) to evaluate the importance of the frequency-versus-time structure of the stimuli for deviance detection. The amplitude-modulation of the communication AM call was produced by the animal itself and the call represents another example of a natural distress call of C. perspicillata. Both vocalisation mimics are artificially generated. They resemble the natural vocalisations in their frequency range and duration but not in their temporal structure.

## 2. Results and Discussion

### Deviance detection in broadband filtered ABRs differs between echolocation and social communication sounds

In experiment 1, an echolocation pulse and a social communication call were presented in the oddball paradigm to investigate differences of deviance detection between both vocalisation types with the following results: ABRs to the echolocation pulse were significantly larger when the stimulus was perceived as a low-probability deviant (Fig. 2a, red ABR) than when it was a high-probability standard (blue ABR). This difference is present across the whole response, however, most prominently it appears in the last peak of the ABR, a slow wave that only becomes visible when the responses are broadband filtered between 0.1 and 2500 Hz. This filtering method is different from the usual narrowband filters between 300-2500 Hz that are used in many ABR studies and that will be discussed later. The strong effect of deviance detection in this late part of the response is in line with previous studies that investigated deviance detection in broadband filtered ABRs with pure tones[7,8]. This confirms that this slow, most likely inferior colliculus-generated[9] wave plays a key role in ABR-based deviance detection. It has been proposed that deviance detection is driven by two mechanisms at the neural level: repetition suppression and deviant enhancement. To disentangle which mechanism underlies the neural responses, the MS control has been suggested[10] (Fig. 1c, top). In this control, the target stimuli (Fig. 1a) are pseudo randomly presented together with multiple other stimuli (here: 8 stimuli; Fig. 1d; Supp. Fig. 1), which makes it impossible for the brain to detect regularities or deviations in the acoustic input. This results in responses that are unaffected by repetition suppression and deviant enhancement. A reduction of response strength to the standard relative to the MS response indicates repetition suppression while a stronger deviant than MS response is evidence for deviant enhancement. The echolocation response that was recorded in the MS control was significantly smaller than the deviant response and not significantly different from the standard response (Fig. 2a, top). This observation shows that the neural mechanism driving deviance detection for echolocation is a deviant-related enhancement of the response (i.e., a prediction error response in the predictive coding framework) and not a repetition suppression effect on the standard response. Possibly, deviant stimuli cause the brainstem neurons to respond more synchronously than standard or MS stimuli, resulting in larger deviant ABR amplitudes. In line with this hypothesis, former studies have demonstrated the importance of synchronisation and phase locking of brainstem neurons for speech[11] and music[12,13] perception in humans. Interestingly, the slow wave of the MS response has an earlier peak and offset latency compared to the deviant and standard ABR. This could indicate that additional neural mechanisms become active and modify the ABRs when the natural acoustic input becomes more complex, as occurs in the MS control compared to the oddball sequence. To further investigate deviance detection in the ABR, we used another common control paradigm, the so-called 50 % control. Here, both target stimuli are presented in a sequence with equal probability of 50 %. The analysis yielded a similar result to the MS control, that is the deviant response being significantly enlarged and no difference between control and standard response (Fig. 2a, bottom). The fact that deviant enhancement and not repetition suppression is driving low-level deviance detection for echolocation calls is interesting since previous studies have suggested repetition suppression to be the dominant mechanism causing deviance detection in subcortical nuclei[4,5]. However, those studies used pure tones to stimulate individual neurons instead of measuring vocalisation-related summed potentials like we did here. It is possible that echolocation pulses evoke stronger deviant responses and less repetition suppression due to their high behavioural relevance compared to simple tone pips.

**Figure 2:**
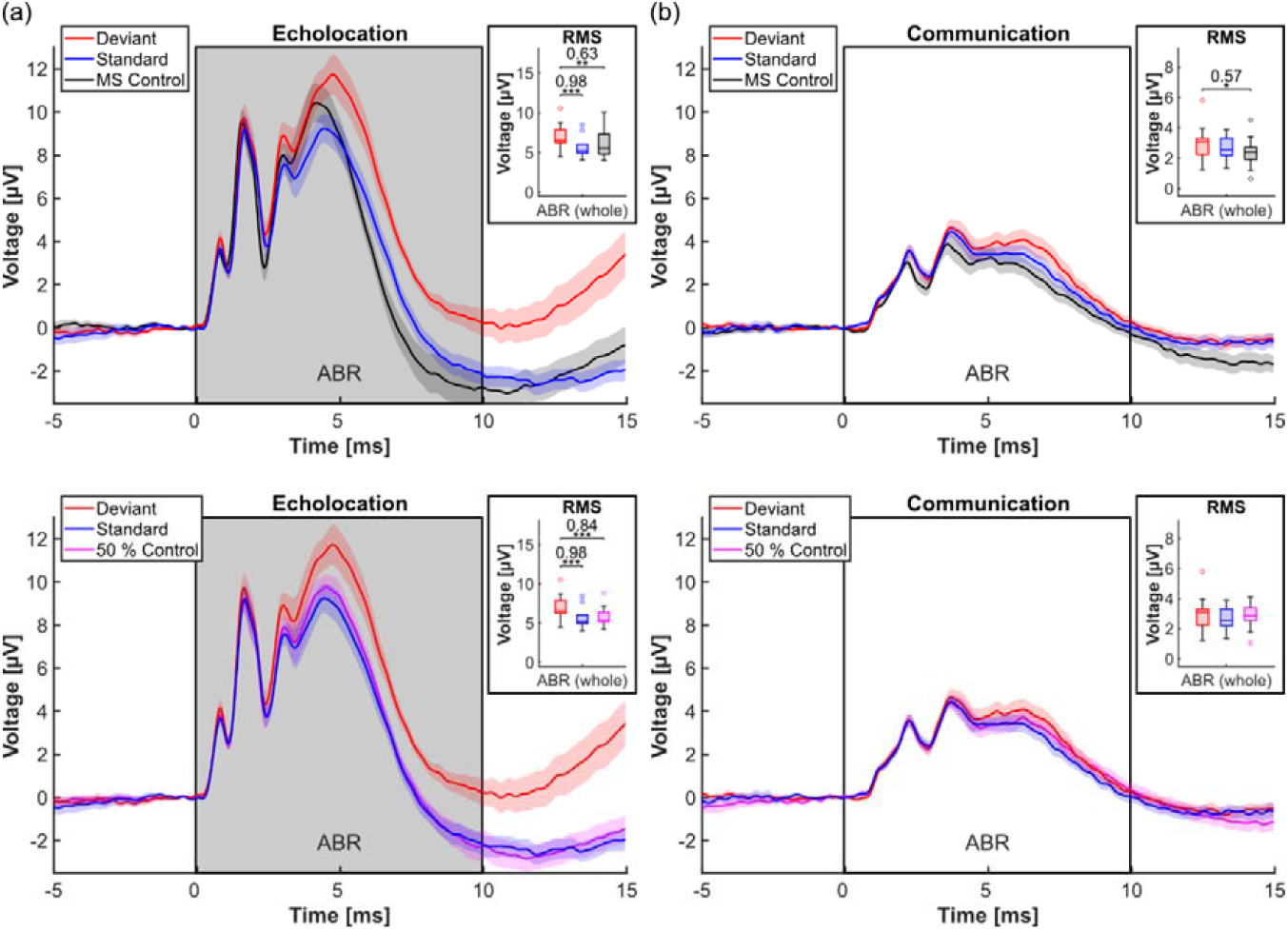
Deviance detection in broadband filtered ABRs differs between echolocation and social communication sounds (n = 13 animals). (a) Grand averages of ABRs to an echolocation pulse presented as deviant (red) and standard (blue) as well as in the MS control (black, top) and the 50 % control (magenta, bottom). The boxes framing the responses represent the time window taken for RMS calculation, covering the whole ABR response (0-10 ms post stimulus onset). The grey colour of the boxes indicates a significant difference between deviant and standard response. Shaded areas around the ABRs depict the standard error of the mean. The inset on the right shows the RMS values calculated for each animal and condition as an estimation of response strength. (b) As in (a) but the stimulus was a social communication call. The white colour of the boxes framing the responses indicates that there was no significant difference between deviant and standard response.

As opposed to echolocation, the social communication sounds did not elicit deviance detection in this experiment (Fig. 2b). While deviant and standard responses were not significantly different from each other, the MS response was attenuated in comparison to the deviant response (Fig. 2b, top). This is surprising since, as explained above, the MS control is expected to generate a baseline response that is affected by neither deviant enhancement nor repetition suppression and hence should be positioned between deviant and standard response. Likely, the attenuation of the MS communication response is the result of the same mechanisms that modified the timing of the MS echolocation response. Those mechanisms seem to get active only when the acoustic input becomes more variable and appear to have complex, nonlinear effects on ABRs. Interestingly, they are restricted to natural stimuli, as a previous study by Wetekam et al.[8] in the same species did not find similar effects in the MS ABRs to pure tones. However, the 50 % control was very similar to the deviant and standard response, confirming that probability encoding did not affect the ABR size to the social communication call (Fig. 2b, bottom).

### For echolocation, deviance detection is measurable very early in narrowband filtered ABRs

To further characterise the effects of deviance detection on the echolocation response, the data were narrowband filtered (bandpass Butterworth, 300-2500 Hz, 4^th^ order) to analyse the fast ABR components in more detail (Fig 3). ABR wave ii/iii as well as wave iv of the deviant response were significantly larger than the respective components of the standard response. Given that wave ii and iii represent neural activity in the cochlear nucleus and superior olivary complex, respectively[14], this finding strongly supports the hypothesis that auditory probability encoding at population level is happening already below the inferior colliculus, as it has been suggested in former studies[8,15]. In fact, effects of novelty detection have recently been described for an even lower auditory structure, the cochlea[16,17]. In these reports, the authors propose that the medial olivocochlear reflex is responsible for those effects by suppressing outer hair cell activity, mediated by feedback from the cortex. Since the ABRs presented here are averaged over many trials, it is possible that similar cortical feedback mechanisms are responsible for the very early effects seen in our ABR data.

**Figure 3:**
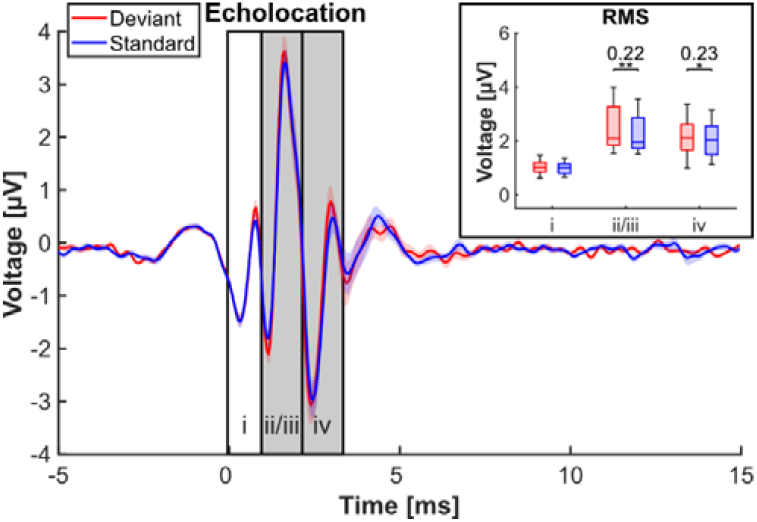
For echolocation, deviance detection is measurable very early in narrowband filtered ABRs (n = 13 animals). Grand averages of ABRs to an echolocation pulse presented as deviant (red) and standard (blue), with a social communication call as context. The boxes framing the responses represent the time window taken for RMS calculation, covering the typical ABR peaks i, ii/iii and iv. The colour of the boxes indicates whether a significant difference between deviant and standard response could be measured (grey: yes, white: no). Shaded areas around the graphs depict the standard error of the mean. The inset on the right shows the RMS values calculated for each animal, condition and time window as an estimation of response strength.

### Behavioural meaning and complex sound features of a stimulus affect deviance detection in broadband filtered ABRs

The second experiment of this paper tackles the question of how low-level deviance detection is affected by individual stimulus parameters and possible behavioural meaning of the stimuli. Therefore, in addition to the previously used echolocation pulse and social communication call (Fig. 1a), an *amplitude-modulated* communication call (another distress vocalisation of *C. perspicillata*) and two artificial vocalisation-mimics that resembled the natural vocalisations in their frequency range and duration but not in their temporal structure (Fig. 1d) served as stimuli. The aim was to assess the relevance of the frequency-versus-time structure of a signal for producing deviance detection in broadband filtered ABRs. In addition, it was tested whether the AM property of a communication call influenced deviance detection, as amplitude modulation appears in natural communication calls[2,18] and could bear additional meaning for the animal. To answer these questions, the five different stimuli were presented to the animals in all possible parings of the oddball paradigm.

When the echolocation pulse or the echolocation mimic served as target stimulus, significantly larger deviant than standard responses could be measured when any communication stimulus was the context (Fig. 4, Supp. Fig. 2 for statistics). Interestingly, deviance detection could also be recorded for the responses to the echolocation pulse when the echolocation mimic was the context, but not vice versa. This indicates that differences in auditory input beyond simple frequency deviations – e.g., the frequency modulation of the echolocation pulse that is absent in the mimic – have a direct influence on subthalamic deviance detection in the bat brain. The fact that this effect is not present in the echolocation mimic responses when the echolocation pulse was context supports the claim that the behavioural meaning of a stimulus plays a key role in low-level population-based deviance detection. Modulatory effects of the behavioural meaning of a stimulus on the strength of deviance detection has previously been known for cortical areas[19], but not for the brainstem. Both natural communication calls – whether amplitude modulated or not – did not reveal significant deviance detection in any oddball combination except when presented with the echolocation mimic. This exception could be due to the very different physical properties of both call-types where the artificial nature of the mimic increases the contrast even further. On the other hand, as in experiment 1, the natural echolocation pulse as context did not cause deviance detection in the responses to either of the natural communication calls. Evidence for differences in the processing of novelty detection between echolocation and communication stimuli in *C. perspicillata* has been reported before[20,21] and is in line with the current data. A possible reason for this phenomenon is the fact that both natural communication calls used in this study are distress calls that the animal emits when it is under physical duress[1,2]. Those distress calls might always elicit the strongest possible neural response in the brains of conspecifics due to the relevance and importance their perception has for the behavioural response, independent of their probability of occurrence. In contrast, ABRs to the communication mimic did reveal strong deviance detection with significantly enlarged deviant responses when the AM communication call, the echolocation pulse or the echolocation mimic was the context. Only when the unmodulated communication call was the context, no significant difference between deviant and standard response could be measured. Together, these results indicate that the AM of the communication stimulus contributed to the differentiation between the true call and an artificial sound while it did not have a significant impact on the distinction between two different natural communication calls at subthalamic level.

**Figure 4:**
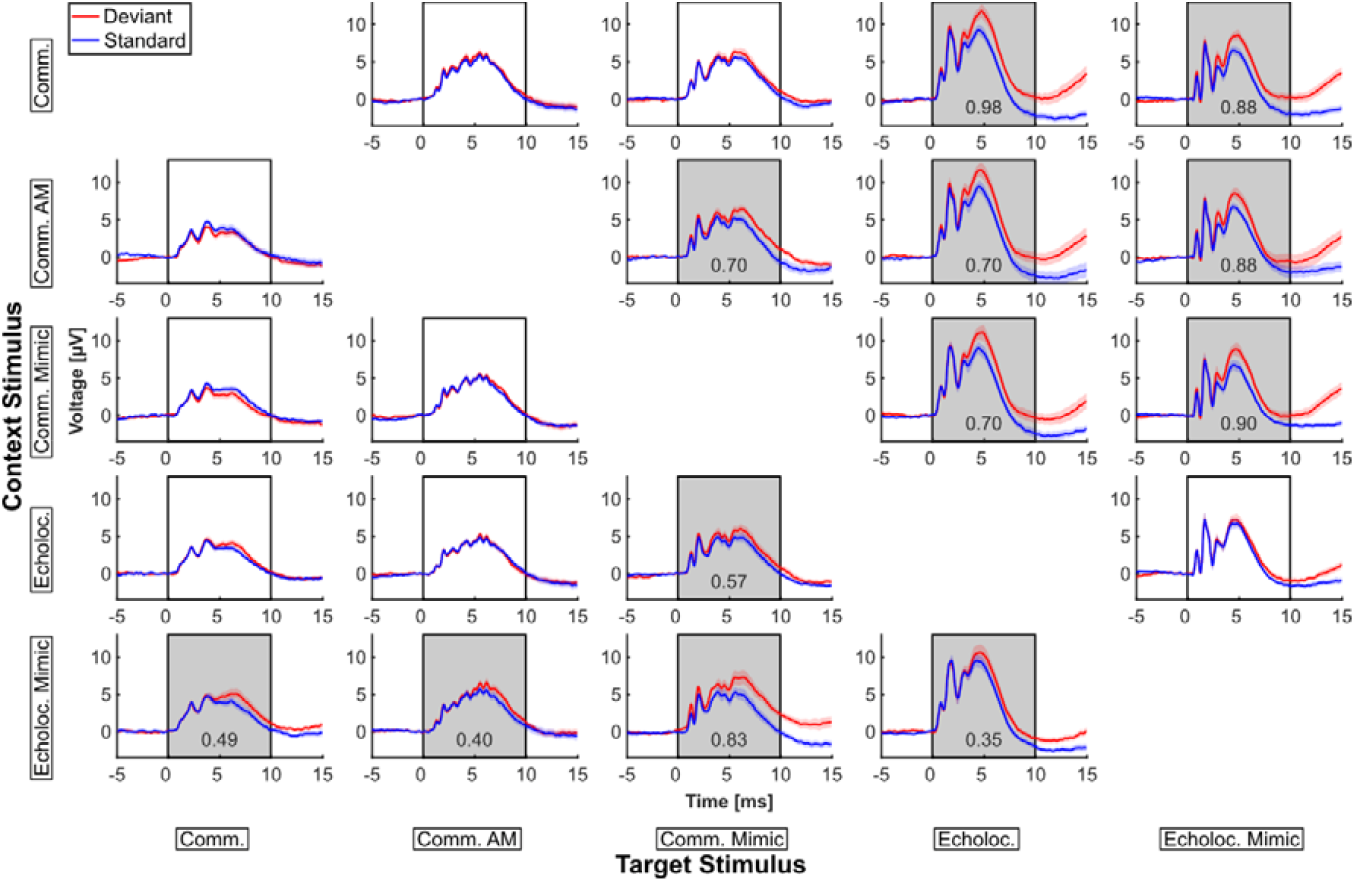
Behavioural meaning and complex sound features of a stimulus affect deviance detection in broadband filtered ABRs (n = 13 animals). All possible parings of the oddball paradigm. Each column contains the recorded responses to the target stimulus of the oddball sequence while each row represents one context stimulus (the second stimulus of the oddball paradigm that served as context for the target stimulus). The stimuli tested were: communication call (Comm.), AM communication call (Comm. AM), communication-mimic (Comm. Mimic), echolocation pulse (Echoloc.) and echolocation mimic (Echoloc. Mimic). Response plots like in Fig. 2. The colour of the boxes indicates whether a significant difference between deviant and standard response could be measured (grey: yes, white: no). If deviant and standard response differed significantly, Cohen’s D is provided as a measure of effect size (number in the grey boxes).

### Conclusion

In this study, noninvasively recorded ABRs revealed that deviance detection responses to vocalisations that have different behavioural meanings for bats – navigation and communication – are processed in a complex and asymmetric way already at the earliest stations of the ascending auditory pathway. In fact, the results show that when considering the population response, subthalamic deviance detection is sensitive to physical (carrier frequency, FM and AM) as well as abstract stimulus features (behavioural meaning of a vocalisation). By this, population-based subthalamic deviance detection showed a higher complexity than what has been reported for cellular SSA of neurons in the same brain areas.

## 3. Material and Methods

### Animals

For the experiments of this study, 13 adult bats (7 males, 6 females) of the species *Carollia perspicillata* from the breeding colony of Goethe University Frankfurt were used. After being caught for the first time, all animals were held separately from the colony until the end of the study. Before every recording session, the animal was anaesthetised by a mixture of ketamine (Ketavet © 10 %, Medistar GmbH Ascheberg, Germany; 7.5 mg per kg bodyweight) and xylazine (Rompun © 2 %, Bayer HealthCare AG, Mohnheim, Germany; 16.5 mg per kg bodyweight) and the anaesthesia was maintained by follow-up injections of the same mixture with reduced volume every 1-1.5 h, for up to 4 h total. A DC-powered heating pad that was attached to the animal holder was used to maintain the animal’s body temperature of 37 °C. Two consecutive recording sessions in the same animal were at least five days apart. This study was approved by the Regierungspräsidium Darmstadt (permits: FR/1010 and FR/2007) and was performed in full compliance with current German laws.

### Stimulation and recording procedure

Custom written MATLAB (MathWorks Inc., US) scripts were used for stimulation and data acquisition. The digital stimulus signal was D/A-converted by a 384 kHz Adi-2 Pro soundcard (RME, Haimhausen, Germany) before it was fed into a HiFi-amplifier (Power Amplifier RB-1050, Rotel, Hongkong, China) and presented to the animal by a Fountek NeoPro 5i Ribbon Tweeter (Fountek Electronics Co.,Ltd, Jiaxing, China). The speaker was positioned 15 cm away from the animal and pointed directly towards the left ear in a 45 ° azimuth angle relative to the head. To ensure a constant distance and angle between ear and speaker, the animal was head-fixed by a mouth-holder. All stimuli were natural vocalisations of *C. perspicillata* or vocalisation mimics with durations between about 2 to 10 ms (Fig 1, Supp. Fig. 1). The social communication call that was used as target tone in experiment 1 (Fig. 1a) is a distress call, a social vocalisation that is emitted by the animal when under physical duress. Like all calls used in this study, it was recorded from a freely behaving bat. The echolocation and communication mimics are noise bursts covering very similar frequency ranges as their natural counterparts. They also resemble the vocalisations in their durations and rise/fall-times, with only the temporal structure of the natural and artificial stimuli being fundamentally different from each other. All stimuli had an intensity of 60 dB SPL and were presented at a rate of 20 Hz, equivalent to a stimulus-onset asynchrony of 50 ms.

The oddball paradigm that was used to study effects of deviance detection in the ABR consisted of 2 sequences of stimuli. In the first sequence, stimulus 1 was presented as standard (high probability, 90 %) and stimulus 2 as deviant (low probability, 10 %). The second sequence was presented consecutively and resembled the first one but with opposite roles of the stimuli, where now stimulus 1 was the deviant and stimulus 2 the standard (Fig. 1b). In total, a sequence contained 1000 stimuli (900 standards, 100 deviants). To characterise the measured deviance detection effects in more detail, two control sequences were used. The first was the MS control[10], presenting the target stimuli (echolocation pulse and social communication call) in a pseudo randomly arranged sequence together with eight additional stimuli, all having a probability of occurrence of 10 %. The other eight stimuli were the two vocalisation mimics, an AM communication (distress) call (Fig. 1d) and 5 other social communications of *C. perspicillata* that are related to different behaviours (Supp. Fig. 1). The MS control is expected to generate responses that are unaffected by any modulatory effects of probability encoding (repetition suppression or deviant enhancement) since the stimuli are perceived neither as deviant nor standard. As a second control, the echolocation pulse and social communication call were presented in another sequence where their probability of occurrence was 50 %, respectively. Like in the oddball paradigm, the sequences of both controls consisted of 1000 stimuli each.

ABRs were differentially recorded by two electrodes – chlorinated silver wires (AG-10T, diameter: 0.25 mm; uninsulated and chlorinated tip of 3 mm) – that were placed subcutaneously at the vertex of the animal’s skull and close to the bulla of the left ear. A ground electrode was clipped to the animal’s right thumb. The recorded responses were hardware filtered (0.1-3000 Hz, 20 dB/decade roll-offs) and amplified by a factor of 20k by a Dagan EX1 differential amplifier (Science Products GmbH, Hofheim, Germany) before they were A/D-converted by the soundcard and sent to the computer. Blocks of 20 consecutive points of the input signal were averaged in order to down-sample the signal to 19.2 kHz.

### Data processing and statistical evaluation

All processing and statistical evaluation of the data was conducted in MATLAB. The recorded ABRs were bandpass filtered by a Butterworth filter (4^th^ order) in two different ways, dependent on the analysis. For the broadband filtered responses, low- and high-cut frequencies of 0.1 Hz and 2500 Hz, respectively, were used, which did remove high frequency noise from the signal but left the ABRs otherwise almost unchanged. On the other hand, the narrowband filter removed frequencies below 300 Hz and above 2500 Hz, abolishing all slow components of the response and allowing a more detailed inspection of the fast ABR waves i-iv. Before averaging, each trial was baseline corrected by calculating the mean voltage in a time window 1 ms pre stimulus onset. This value was subtracted from the whole trail resulting in a pre-stimulus activity of 0 μV. Subsequently, the averaging procedure was restricted to those deviant responses that followed a standard response and, vice versa, those standard responses that preceded a deviant response. This method allows to use the same number of trials to calculate the deviant and standard average of each animal (here between 89 and 92 trials) and, at the same time, maximises the effects of deviance detection in the responses[8]. In the case of the 50 % control, the same number of trials that was used to calculate the deviant and standard average for a given animal was used to randomly choose trials out of the 500 available responses to each stimulus. The MS average was calculated based on all responses to a given stimulus in the MS sequence (between 85 and 110 trials; mean difference to oddball responses: + 2.4 trials). All responses were corrected for the sound-travelling delay caused by the distance between speaker and ear. In each graph, the time point of 0 ms represents the moment when the sound reached the bat’s ear.

To evaluate the response strength of each ABR, time windows were defined within which the response’s RMS value was calculated. This has been done successfully in the same species before[8,22]. For all broadband filtered responses, this time window spanned from 0 ms to 10 ms, covering the whole ABR with all its fast and slow components. The detailed wave-by-wave analysis of the narrowband filtered responses was done using three consecutive time windows, each containing a different component of the ABR. Those windows had borders of 0 ms – 1 ms (wave i), 1 ms – 2.2 ms (wave ii/iii) and 2.2 ms – 3.4 ms (wave iv) which are similar to previously reported ABR-wave latencies of other bat species[23–25].

To compare response strengths between conditions with each other, paired one-tailed t-tests (deviant vs. standard responses) and repeated measure ANOVAs (deviant vs. standard vs. control responses) with subsequent Bonferroni-corrected post-hoc tests were used to evaluate differences between the calculated RMS values. Additionally, the effect size measure Cohen’s D was calculated for all significant comparisons which allows an estimation of strength of the measured deviance detection effects[26].

## Acknowledgements

We would like to thank Dr. Mirjam Knörnschild for providing us with some of the vocalisations of *Carollia perspicillata* that were used in this study. The study was funded by the Deutsche Forschungsgemeinschaft (KO 987/14-1).

## Author contributions

Study design: Johannes Wetekam, Julio Hechavarría, Luciana López-Jury, Eugenia González-Palomares, Manfred Kössl

Experiments, analysis, and original draft of the manuscript: Johannes Wetekam

Review and editing of the original draft: Johannes Wetekam, Julio Hechavarría, Luciana López-Jury, Eugenia González-Palomares, Manfred Kössl

## Conflicts of interest

The authors declare no competing interests.

## Abbreviations

ABR: Auditory brainstem response
AM: Amplitude modulation
ANOVA: Analysis of variance
FM: Frequency modulation
MMN: Mismatch negativity
MS: Many-standards
RMS: Root Mean Square
SSA: Stimulus-specific adaptation

## Notes

### Competing Interest Statement

The authors have declared no competing interest.

