## Supplementary Figures for "Deviance detection in subthalamic neural population responses to natural stimuli in bats"

### Supplementary material

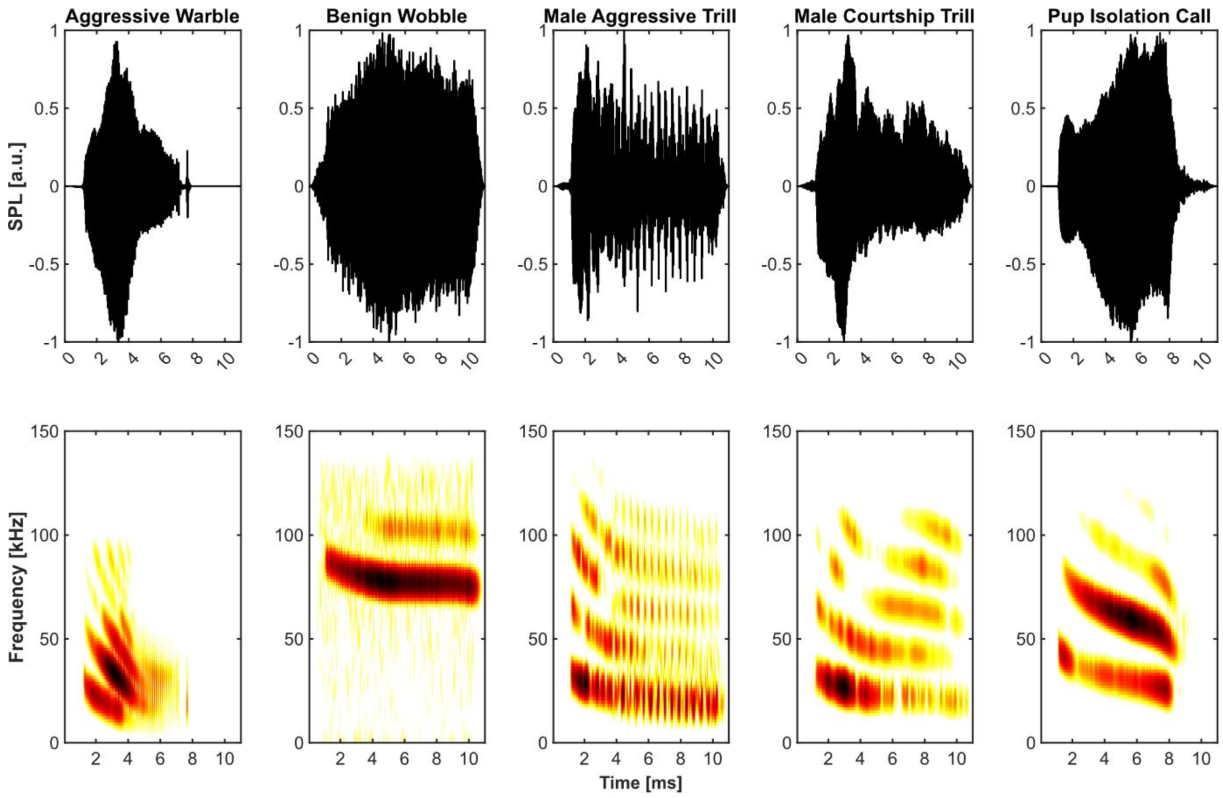


*Supplementary Figure 1: Additional stimuli used in the Many-Standards control. Five additional vocalisations of C. perspicillata that were presented together with the stimuli in Fig. 1 in the Many-Standards control.*

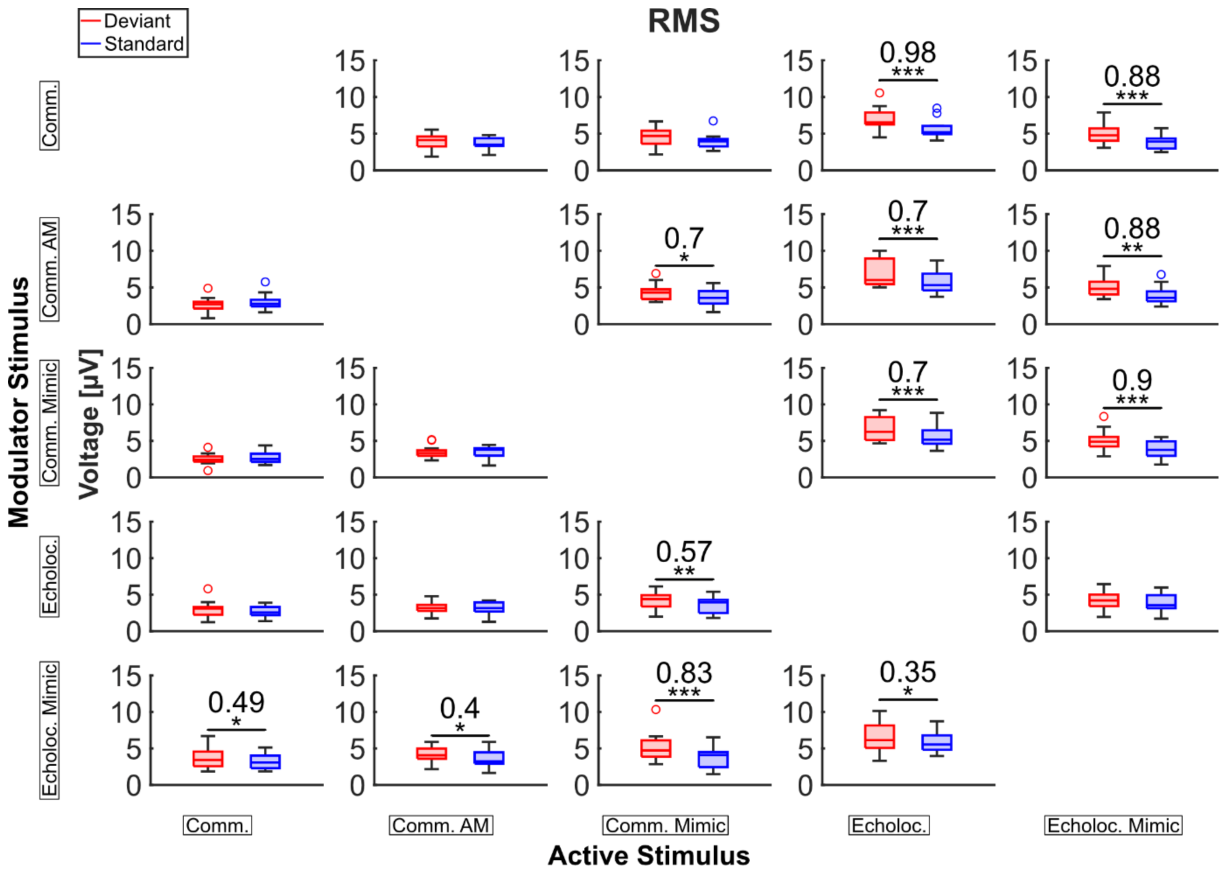


*Supplementary Figure 2: RMS values for each stimulus combination and condition in the oddball paradigms of experiment 2.*
